# Genome-wide identification and characterisation of HOT regions in the human genome

**DOI:** 10.1101/036152

**Authors:** Hao Li, Feng Liu, Chao Ren, Xiaochen Bo, Wenjie Shu

## Abstract

HOT (high-occupancy target) regions, which are bound by a surprisingly large number of transcription factors, are considered to be among the most intriguing findings of recent years. An improved understanding of the roles that HOT regions play in biology would be afforded by knowing the constellation of factors that constitute these domains and by identifying HOT regions across the spectrum of human cell types. We characterised and validated HOT regions in embryonic stem cells (ESCs) and produced a catalogue of HOT regions in a broad range of human cell types. We found that HOT regions are associated with genes that control and define the developmental processes of the respective cell and tissue types. We also showed evidence of the developmental persistence of HOT regions at primitive enhancers and demonstrate unique signatures of HOT regions that distinguish them from typical enhancers and super-enhancers. Finally, we performed an epigenetic analysis to reveal the dynamic epigenetic regulation of HOT regions upon H1 differentiation. Taken together, our results provide a resource for the functional exploration of HOT regions and extend our understanding of the key roles of HOT regions in development and differentiation.

## Introduction

Recent studies in *Caenorhabditis elegans* [1, 2], *Drosophila melanogaster* [3–7], and humans [8–10] have identified a class of mysterious genomic regions that are bound by a surprisingly large number of transcription factors (TFs) that are often functionally unrelated and lack their consensus binding motifs. These regions are called HOT (high-occupancy target) regions or “hotspots”. In *C. elegans*, 22 different TFs were used to identify 304 HOT regions bound to 15 or more TFs [1]. Using the binding profiles of 41 different TFs, nearly 2,000 HOT regions were identified in *D. melanogaster*, and each is bound by an average of 10 different TFs [5]. Many regions that are bound by dozens of TFs were also identified in a small number of human cells [9, 10]. The broad presence of these regions in metazoan genomes suggests that they might reflect a general property of regulatory genomes. However, how hundreds of TFs coordinate clustered binding to regulatory DNA to form HOT regions across cell types and tissues is still unclear. Furthermore, the function of HOT regions in gene regulation remains unclear [11, 12], and their proposed roles include functioning as mediators of ubiquitously expressed genes [1], sinks or buffers for sequestering excess TFs [4], insulators [5], DNA origins of replication [5], and patterned developmental enhancers [7]. In addition, the effects of these regions on human diseases and cancer remain unknown. Thus, it is important to systematically analyse HOT regions in a large variety of cell types and tissues and to further understand their functional roles in the control of specific gene expression programs.

Resolving these challenges requires knowledge of the ensemble of all TF bindings in a cell. However, even predicting where a single TF binds in the genome has proven challenging. Computational motif discovery in regulatory DNA is a commonly used strategy for identifying candidate TF binding sites (TFBSs) for TFs with known binding motifs, which are represented as position weight matrices (PWMs). Numerous algorithms have been developed for discovering motifs, such as FIMO (Find Individual Motif Occurrences) [13] and HOMER (Hypergeometric Optimization of Motif EnRichment) [14]. It has been reported that TFBSs tend to be DNase I hypersensitive, and only a fraction of the human genome is accessible for TF binding [15]. Remarkably, HOT regions correlate with decreased nucleosome density and increased nucleosome turnover and are primarily associated with open chromatin [1, 5, 6]. DNase I hypersensitive sites (DHSs) in chromatin have been used extensively to mark regulatory DNA and to map active *cis*-regulatory elements in diverse organisms [16–18]. Recent advances in Next-Generation Sequencing (NGS) technologies have enabled genome-wide mapping of DHSs in mammalian cells [19–21], laying the foundations for comprehensive catalogues of human regulatory DNA regions. Thus, DHSs, combined with motif discovery algorithms, could be used in a very powerful approach for identifying a large repertoire of TFs in diverse cell and tissue types with high precision. This approach is likely to be widely applicable for investigating cooperativity among TFs that control diverse biological processes.

Here, we have developed a computational method for the genome-wide mapping of HOT regions in human genome. We have characterised and validated HOT regions in embryonic stem cells (ESCs). Additionally, we have created a catalogue of HOT regions for 154 different human cell and tissue types and have shown that these regions are associated with genes encoding cell-type-specific TFs and other components that play important roles in cell-type-specific developmental processes. We have shown evidence for the developmental persistence of HOT regions at primitive enhancers and have demonstrated unique signatures of HOT regions that distinguish them from typical enhancers and super-enhancers. Importantly, our epigenetic analysis revealed a preliminary view of dynamic epigenetic regulation of HOT regions upon cell differentiation.

## Results

### Identification and validation of HOT regions

Recently, we used Gaussian kernel density estimation across the binding profiles of 542 TFs to identify TFBS-clustered regions and defined a “TFBS complexity” score based on the number and proximity of contributing TFBSs for each TFBS-clustered region [22]. A preliminary inspection of these regions with different TFBS complexity revealed an unusual feature: Although the vast majority (~90%) of the TFBS-clustered regions exhibited only low TFBS complexity, a small portion of the TFBS-clustered regions exhibited much higher TFBS complexity scores (Fig. S1A and Table S1). The former were called LOT (low-occupancy target) regions, whereas the latter were called HOT (high-occupancy target) regions.

HOT regions were initially defined as regions with high occupancy of TFs and were identified by the binding peaks of many TFs using ChIP-seq data in previous reports [1–10]. However, we defined HOT regions by the colocalisation of a large number of TF motif binding sites and identified them using the TF motif scanning method iFORM [23] on DHSs with PWMs for 542 TFs. To validate our identified HOT regions, we compared them with the experimental HOT regions that were defined by the ENCODE Consortium, which assessed more than 100 TFs from approximately 500 ChIP-seq experiments in more than 70 cell types [24, 25]. We performed a GSC (genome structure correction) analysis between our HOT regions and the experimental HOT regions in five cell types, including H1-hESC, K562, Hep-G2, HeLa-S3, and GM12878 cells (Fig. 1A and S1B). The GSC results indicated that our HOT regions were significantly enriched and depleted compared to the experimental HOT and LOT regions, respectively. In addition, our LOT regions were significantly enriched and depleted compared to the experimental LOT and HOT regions, respectively. Furthermore, we used the experimental HOT and LOT regions to validate our HOT regions with ROC (receiver operating characteristic) curves and the corresponding area under the curve (AUC) in each cell type. Our results demonstrated that our HOT regions have powerful discrimination ability between the experimental HOT and LOT regions (Fig. S1C).

**Figure 1.**
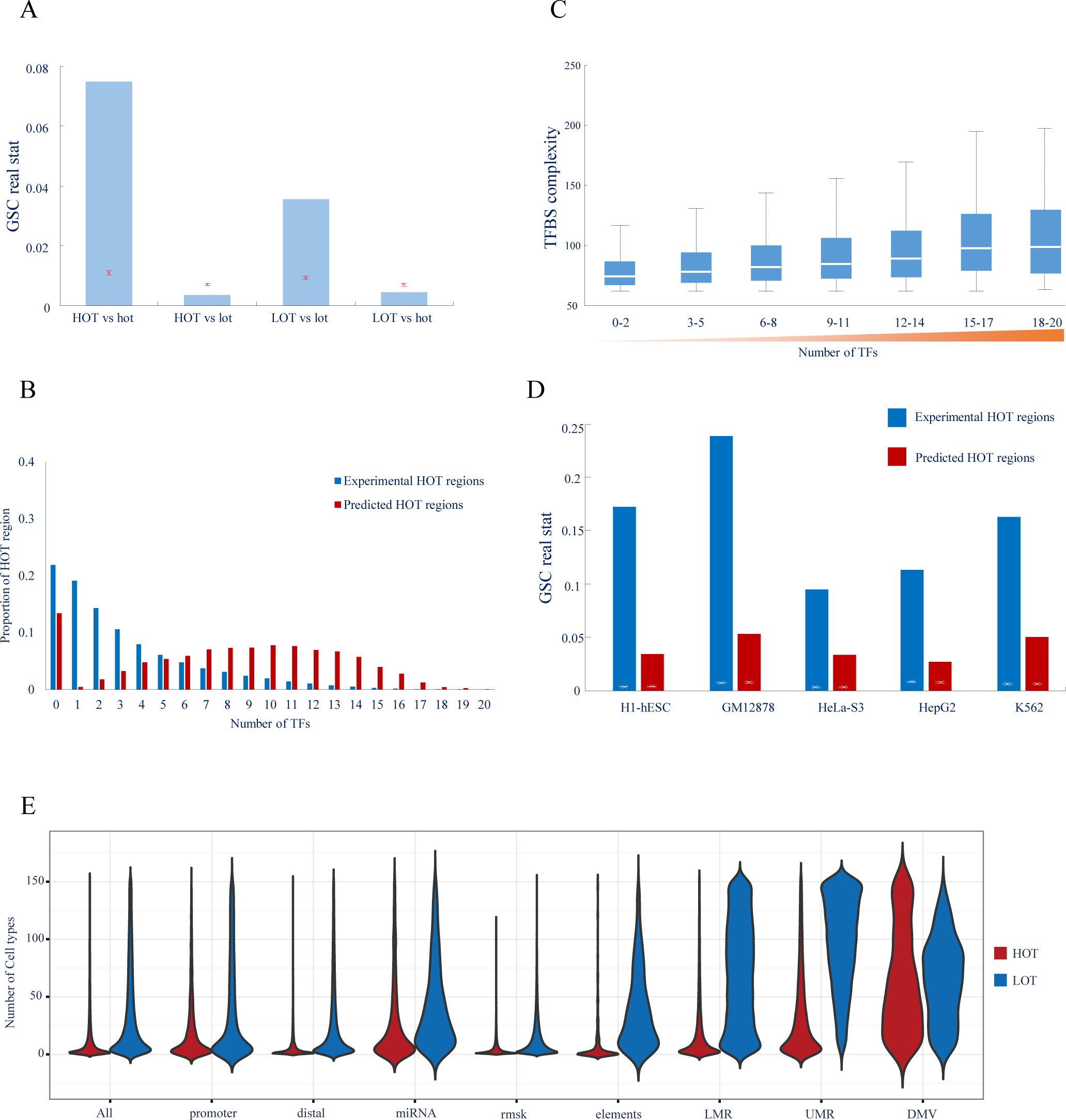
Identification and validation of HOT regions. (A) Error bar showing the GSC results for HOT/LOT regions versus experimental HOT/LOT regions. Red lines indicate the mean and normalised SD of 10,000 bootstrap samples; the blue bar indicates the real statistics. (B) The proportions of HOT regions and experimental HOT regions containing different numbers of ChIP-seq peaks corresponding to TFs in H1 cells. (C) The distributions of TF complexity of HOT regions containing different numbers of ChIP-seq peaks corresponding to TFs in H1 cells. (D) Error bar showing the GSC results of HOT regions versus motifless binding peaks. White lines indicate the mean and normalised SD of 10,000 bootstrap samples; blue and red bars indicate the real statistics of experimental HOT regions and our predicted HOT regions, respectively. (E) Distributions of the number of cell types, from 1 to 154 (*y* axis), in which HOT (red) and LOT (blue) regions in each of nine classes (*x* axis) are observed. The width of each shape at a given *y* value shows the relative frequency of regions present in that number of cell types. See also Figure S1-S3 and Table S1-S5.

To further verify whether TFs indeed bound within the HOT regions, we counted the occurrence rates of peaks in the ChIP-seq data that corresponded to diverse TFs that were located within our HOT regions and the experimental HOT regions. We found that the number of TFs that colocalised within our HOT regions (median = 9 and mean = 8.18 in H1 cells) was much greater than the number of TFs that colocalised within the experimental HOT regions (median = 2 and mean = 3.14 in H1 cells) (Fig. 1B and S1D). Our results suggest that our HOT regions are strongly skewed relative to the experimental HOT regions toward occupancy by a large number of transcription factors identified via ChIP-seq experiments by the ENCODE Consortium. Additionally, with the increase in the TFBS complexity of our HOT regions, the number of TFs that colocalised within our HOT regions also increased (Fig. 1C and S1E).

Previous studies have revealed that some ChIP-seq binding peaks of TFs do not contain the DNA sequence motifs of the corresponding TFs; these peaks are designated motifless binding peaks of the TFs [24, 25]. We explored the relationship between the motifless binding peaks of all TFs and our identified HOT regions. We identified 62,764, 87,582, 129,795, 47,384, and 92,592 motifless binding peaks in H1-hESC, K562, Hep-G2, HeLa-S3, and GM12878 cells, respectively. We compared these motifless binding peaks with the HOT regions that we identified within TF ChIP-seq binding peaks for each cell line. We determined that the proportion of the motifless binding peaks intersecting with the experimental HOT regions (average 25%) was larger than that of the motifless binding peaks intersecting with our HOT regions (average 17%) (Fig. S1F). However, the proportion of motifless HOT regions in our HOT regions was much larger than that of motifless HOT regions in the experimental HOT regions (36% vs 20%, on average) (Fig. S1G). This result reflects the much smaller number and longer length of our HOT regions, Furthermore, GSC analysis demonstrated that the statistical z-scores of the intersections of the motifless binding peaks with our HOT regions and the experimental HOT regions were greater than 57 (corresponding to a *p*-value < 2.2×10^−308^) and 220 (corresponding to a *p*-value < 2.2×10^−308^), respectively (Fig. 1D and Table S2). Our results indicate that although our HOT regions are essentially different from the experimental HOT regions, motifless HOT regions represent a substantial fraction of both our HOT regions and the experimental HOT regions.

Taken together, these findings strongly validate our identified HOT regions. Additionally, the TFBS complexity adequately represented the colocalisation of multiple TFs within our HOT regions. Furthermore, our results clearly clarified the discrimination between our HOT regions and experimental HOT regions that have been reported in the literature [24, 25].

### General characterisation of HOT regions

Using a uniform processing pipeline, we created a catalogue of 59,986 distinct HOT regions across 154 cells/tissues studied under the ENCODE Project [24]. Collectively, these HOT regions span 18.8% of the genome (Table S3). To assess the rate of discovery of new HOT regions, we performed a saturation analysis as described in a previous study [24] and predicted saturation at approximately 107,184 HOT regions, suggesting that we have discovered more than half (59,986 out of 107,184, 56.0%) of the estimated total number of HOT regions (Fig. S2A). An additional location analysis of these 59,986 HOT regions demonstrated that HOT regions were more likely localised to genic regions (intron and exon) and less likely localised to intergenic regions compared with LOT regions (Fig. S2B). Furthermore, HOT regions are typically much more cell-selective than LOT regions (Fig. 1E, 1^st^ column). Promoter proximal HOT regions typically exhibit high accessibility across cell types, with the average proximal HOT region detected in 21 cell types; however, distal HOT regions are largely cell selective, with the average distal HOT region detected in 7 cell types (Fig. 1E, 2^nd^ and 3^rd^ columns).

We further characterised these HOT regions using multiple data types and showed that they are enriched for active histone markers and depleted for repressive histone markers (Fig. S3A), they are highly transcribed (Fig. S3B), they extensively overlap with the transcriptional regulators that control cell development and differentiation (Fig. S3C, S3F), they exhibit distinct sequence signatures (Fig. S3D), and their neighbouring genes illustrate functional enrichment linked to the developmental processes of the respective cell and tissue types (Fig. S3E, Tables S4-S5).

### Associations with functional regulatory elements

To gain understanding of the functional roles of HOT regions, it would be valuable to explore their associations with previously validated regulatory elements. First, we explored the extent to which HOT regions associate with microRNAs, which comprise a major class of regulatory molecules and have been extensively studied, resulting in the consensus annotation of hundreds of conserved microRNA genes [26]. Of 2,633 annotated microRNA transcriptional start sites (TSSs), 1,667 (63%) coincide with a HOT region. The accessibility of HOT regions at microRNA promoters was highly promiscuous compared with GENCODE TSSs (Fig. 1E, 4^th^ column) and showed cell lineage organisation, paralleling the known regulatory roles of well-annotated lineage-specific microRNAs (Fig. 2A). Next, we investigated the association between HOT regions and transposon sequences. A surprising number of these sequences contain highly regulated HOT regions (Fig. 1E, 5^th^ column, and Table S6), which is compatible with the cell type-specific transcription of repetitive elements detected using ENCODE RNA sequencing data [27]. The examples shown in Fig. 2B also illustrate the strong cell-selectivity of chromatin accessibility observed for each major class of repeats. Furthermore, we compared HOT regions with an extensive compilation of 373 experimentally validated distal, non-promoter *cis*-regulatory elements, such as insulators, locus control regions (LCRs), transcription initiation platforms (TIPs), and more. This analysis revealed that the overwhelming majority (76%) of these elements are encompassed within HOT regions (Table S7) and typically show strong cell selectivity (Fig. 1E, 6^th^ column, Fig. 2C and S4A). Finally, we explored the extent to which HOT regions associate with different classes of DNA methylation depleted regions, including low methylation regions (LMRs), unmethylated regions (UMRs), and DNA methylation valleys (DMVs). These DNA methylation-depleted regions have been reported to function as *cis* regulatory elements that are strongly associated with transcription factor genes and developmental genes [28, 29]. Our GSC analysis demonstrated that LMRs, UMRs and DMVs were highly enriched within HOT regions (Fig. S4B-D) and typically showed strong cell selectivity (Fig. 1E, 7-9th column and Fig. 2D). Together, our results suggested that HOT regions are highly associated with the functional regulatory elements that play key developmental roles in a manner that is typically cell-type-specific.

**Figure 2.**
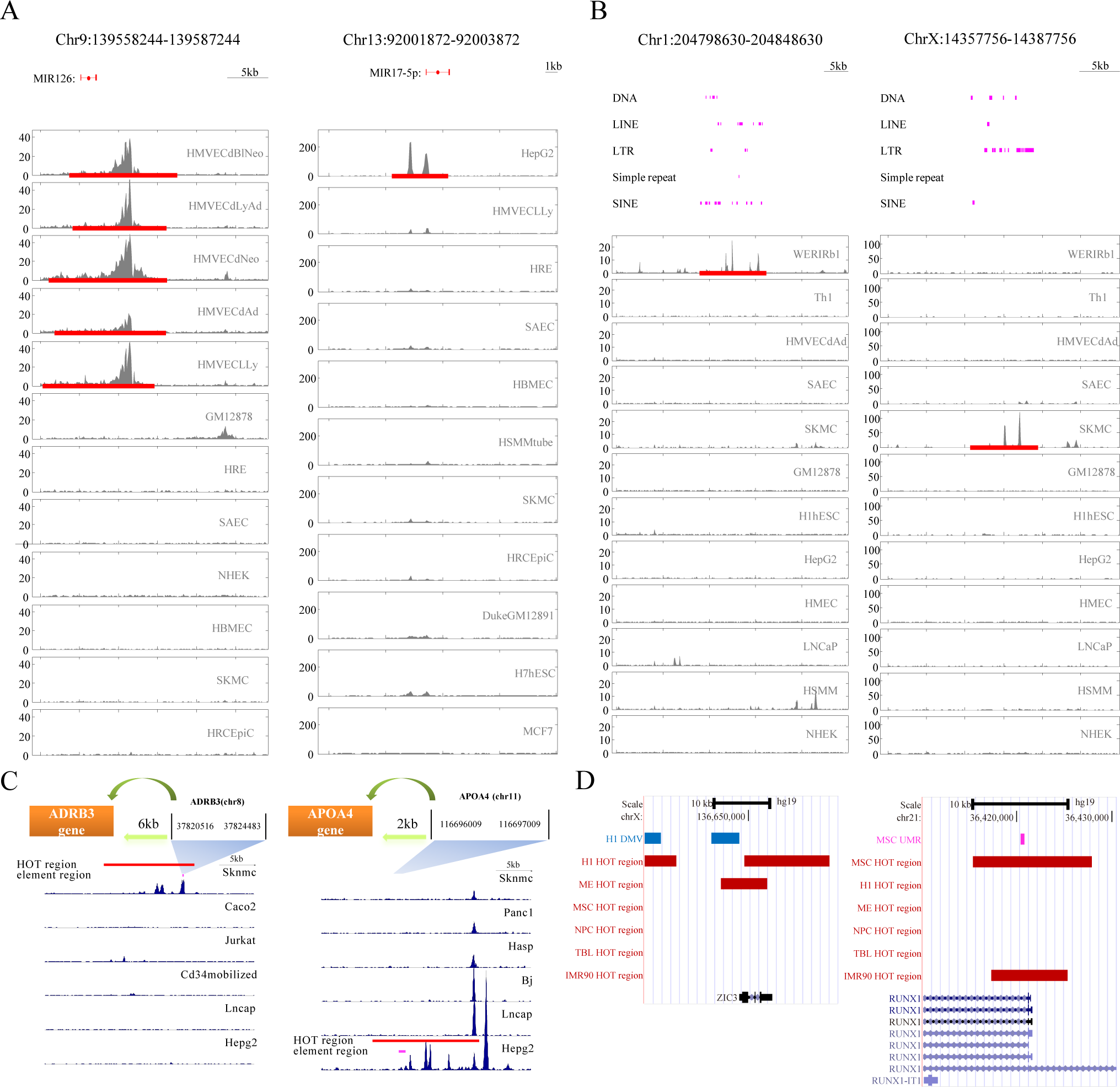
Association of HOT regions with functional elements. (A–B) Examples of HOT regions (red line) overlapping microRNA (A) and repetitive elements (B). Peaks are observed in cell types consistent with known functions of the microRNAs and repetitive elements (pink line). (C) Examples of known cell-selective experimentally validated distal, non-promoter *cis*-regulatory elements. Shown above each set of DNaseI data are schematics displaying HOT regions relative to the genes they control. (D) Examples of colocalisation of DMV (blue) and UMR (pink) with HOT regions (red) in a cell-type-specific manner. See also Figure S4 and Tables S6–S7.

### HOT regions at embryonic enhancers

As HOT regions drive genes that control and define cell development, it is reasonable to surmise that in definitive cells, HOT regions could be persistently associated with enhancers that are active during early development. We compiled 882 early developmental enhancers that were identified through a comparative genome analysis and experimental validation of *in vivo* enhancer activity in transgenic mice [30]. Each of these enhancers displayed reproducible tissue-staining patterns in one or more embryonic tissues at embryonic day 11.5 (Fig. 3A). Of these 882 non-promoter human enhancers, GSC analysis demonstrated that a surprising proportion (308/882, 35%, z-score = 14.9, corresponds to a *p*-value < 6.5 × 10^−49^) occurred within HOT regions in at least one definitive human cell type. To quantify the tissue activity spectra of these embryonic enhancers, we systematically examined their lacZ expression patterns in transgenic mice and related these patterns to HOT region patterning at the same elements across different definitive cell types (Fig. 3B). For example, an enhancer that is selectively active in embryonic forebrain tissue (Fig. 3A, 1^st^ column) was selectively found in HOT regions within cells derived from human ESCs (Fig. 3B, 1^st^ column), and an enhancer that is selectively active in embryonic blood vessels (Fig. 3A, 2^nd^ column) was selectively found in HOT regions within endothelial cells (Fig. 3B, 2^nd^ column). In contrast, an enhancer with extremely broad tissue activity (Fig. 3A, 7^th^ column) was found in HOT regions in nearly all definitive cell types (Fig. 3B, 7^th^ column). These findings were further confirmed across the spectrum of enhancers (Figs. 3C-D). A total of 62.5% of enhancers that are active in embryonic blood vessels were found in HOT regions in endothelial cells, whereas only 26.3% of all other embryonic enhancers were found in endothelial HOT regions. Similarly, 59.3% of enhancers that are active in embryonic heart tissue were found in HOT regions within cells derived from human heart and great vessel structures, whereas only 30.7% of all other embryonic enhancers were found in HOT regions in these cell types.

**Figure 3.**
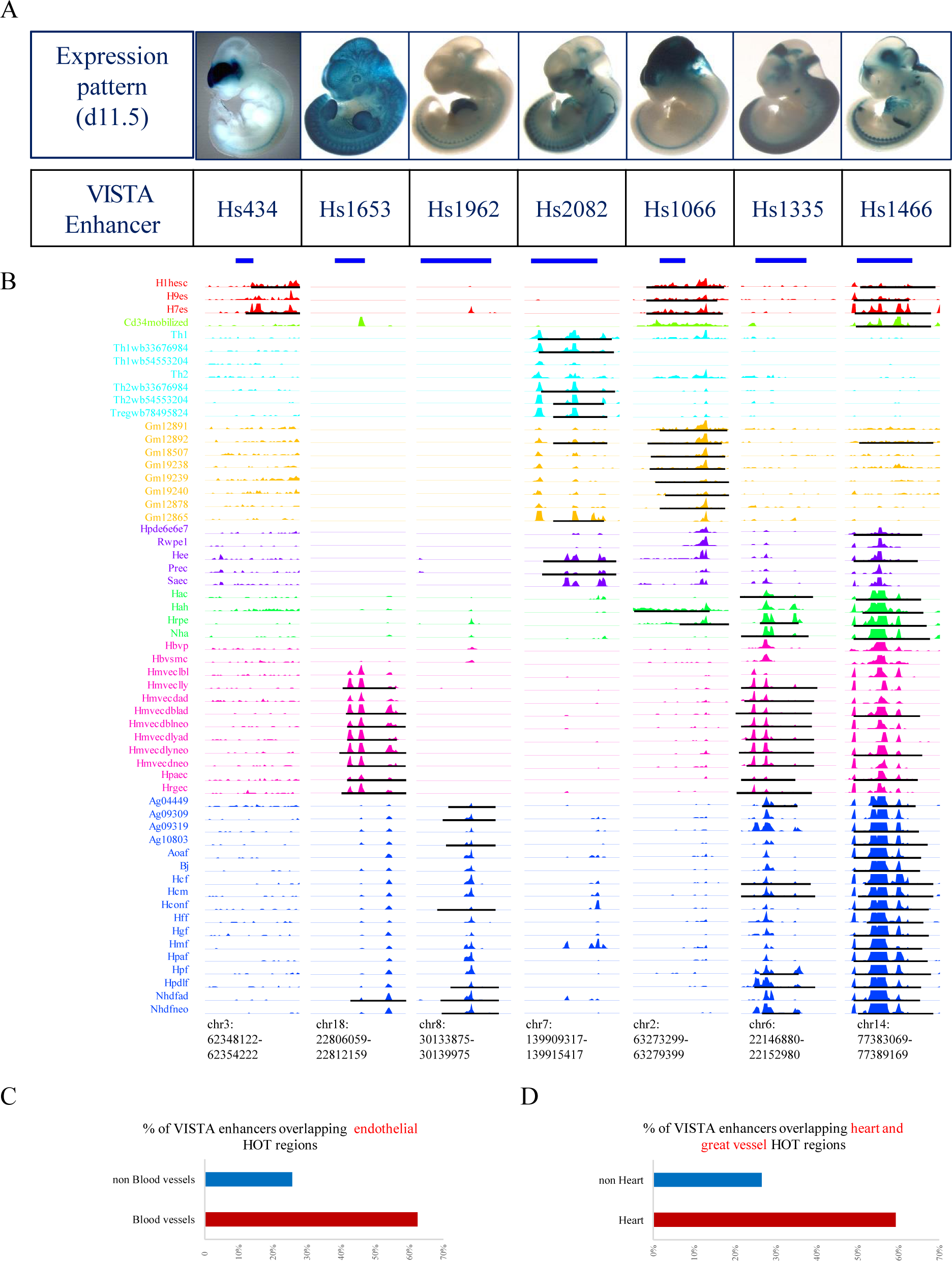
Developmental persistence of HOT regions at embryonic enhancers. (A) Mouse day 11.5 embryonic tissue activity (blue lacZ staining) of seven representative transgenic human enhancer elements from the VISTA database. Shown below each image are the enhancer ID and numbers of individual embryos with enhancer activity (staining) in the indicated anatomical structure. (B) DNaseI hypersensitivity at seven enhancer elements corresponding to (A) across 57 definitive cell types. Note the relationship between the anatomical staining patterns in (A) and the cellular restriction (or lack thereof) of DNaseI hypersensitivity. (C-D) Persistence of HOT regions at embryonic enhancers. (C) Percentage of validated embryonic enhancers from the VISTA database with blood vessel staining (“Blood vessels”) and without blood vessel staining (“NOT Blood vessels”) that overlap a HOT region in any human endothelial cell type. (D) Percentage of validated embryonic enhancers from the VISTA database with heart staining (“Heart”) and without heart staining (“NOT Heart”) that overlap a HOT region in any human paraxial mesoderm cell type.

### Distal HOT regions in super-enhancers

Recently, Richard A. Young and his colleagues identified an unusual class of enhancer domains, called super-enhancers, that drive the high-level expression of genes that control and define cell identity and disease [31–33]. To elucidate the relationship between HOT regions and super-enhancers, we assessed the HOT regions and super-enhancers from the same 14 cell types. We performed a GSC analysis between distal HOT regions and super-enhancers in these cell types and found that HOT regions were significantly enriched with super-enhancers (Fig. 4A). To determine whether HOT regions might cooperate with super-enhancers to regulate cell type-specific gene regulation, we performed a colocalisation analysis of these two types of regions in 14-by-14 cell line combinations, as previously described [34, 35] (Fig. 4B). The diagonally matched cell line enrichment values (> 1.00 for all comparisons) were much larger than the off-diagonal mismatched cell line values (< 1.00 for all comparisons), indicating that cell type-specific HOT regions tended to strongly colocalise with super-enhancers that function in the corresponding cell types. Furthermore, we compared the densities of chromatin markers, TFs, and RNA polymerase II between HOT regions, enhancers, and super-enhancers. All of these elements exhibited similar DNase I hypersensitivity. As expected, enhancer markers, such as H3K27ac, H3K4me1, and P300, were significantly enriched within enhancers and super-enhancers compared to HOT regions. In addition, RNA Pol II was significantly enriched within enhancers and super-enhancers compared to HOT regions. Notably, HOT regions demonstrated simultaneous significant enrichment of the bivalent markers H3K4me3 and H3K27me3, whereas enhancers and super-enhancers showed both enrichment of H3K4me3 and depletion of H3K27me3 compared to the background genome (Fig. 4C). A much higher proportion of HOT regions (28%) were marked with both H3K4me3 and H3K27me3, whereas only 2% of super-enhancers were marked with both H3K4me3 and H3K27me3. Finally, we characterised super-enhancer-associated and HOT region-associated genes by gene ontology (GO) analysis. Our results revealed that super-enhancer-associated genes are linked to biological processes that largely define the identities of the respective cell and tissue types, which is highly consistent with the results of a previous study [31]. However, HOT region-associated genes are linked to biological processes that largely define the development and differentiation of the respective cell and tissue types (Fig. S3E and Fig. S5).

**Figure 4.**
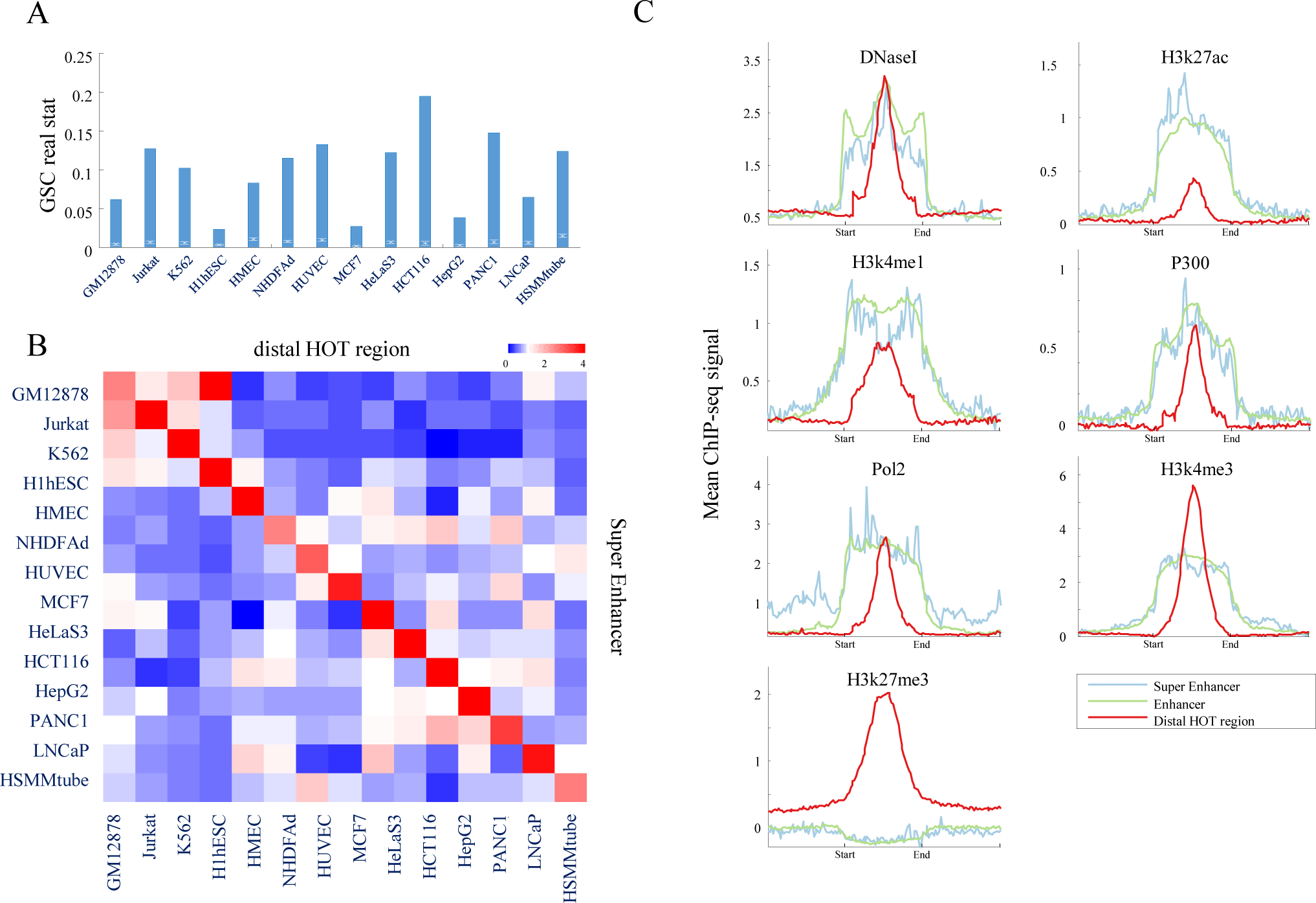
Association of distal HOT regions with super-enhancers. (A) Error bar showing the GSC results between distal HOT regions and super enhancers. Red lines indicate the mean and normalised SD of 10,000 bootstrap samples; blue bar indicates the real statistics. (B) Distal HOT regions colocalise with super-enhancers in a cell type-specific manner. Cell type-specific super-enhancers (*y*-axis) are mapped relative to cell-specific distal HOT regions (*x*-axis) in 14 different cell types. (C) ChIP-seq binding profiles of super-enhancers, enhancers, and distal HOT regions for the indicated DNaseI and enhancer-relevant markers, including transcription factors, transcriptional cofactors, chromatin regulators, and RNA polymerase II in ESCs. See also Figure S5.

### Dynamics of HOT regions during H1 differentiation

To preliminarily explore the dynamic changes in HOT regions upon development and differentiation, we examined the HOT regions during the differentiation of the hESC line H1 to mesendoderm (ME), neural progenitor cells (NPC), trophoblast-like cells (TBL), and mesenchymal stem cells (MSC). We first identified 5,410, 5,602, 4,250, 3,408, and 6,719 HOT regions in H1, ME, NPC, TBL, and MSC, respectively, as well as 4,002 HOT regions in a control for terminally differentiated cells (IMR90).

We first examined the dynamic changes in HOT regions and correlated these dynamic HOT regions with their associated gene expression. During H1 differentiation, we defined “gained” HOT regions as those belonging to H1-derived cell types but not occurring in H1 cells (Fig. 5A). We identified 3,264, 1,784, 2,133 and 4,452 “gained” HOT regions in ME, NPC, TBL, and MSC, respectively. More than half of the HOT regions were “gained” in H1-derived cells upon H1 differentiation, except in NPC (Fig. 5A). We observed that the genes associated with “gained” HOT regions in H1-derived cells were highly expressed compared to those of H1 cells upon H1 differentiation (Fig. 5B, p-value < 1.38×10^−5^, Kolmogorov-Smirnov test). Furthermore, the genes associated with “gained” HOT regions played key roles in development-related biological processes in the respective H1-derived cells in a cell-type-specific manner (Fig. 5C and Table S8).

**Figure 5.**
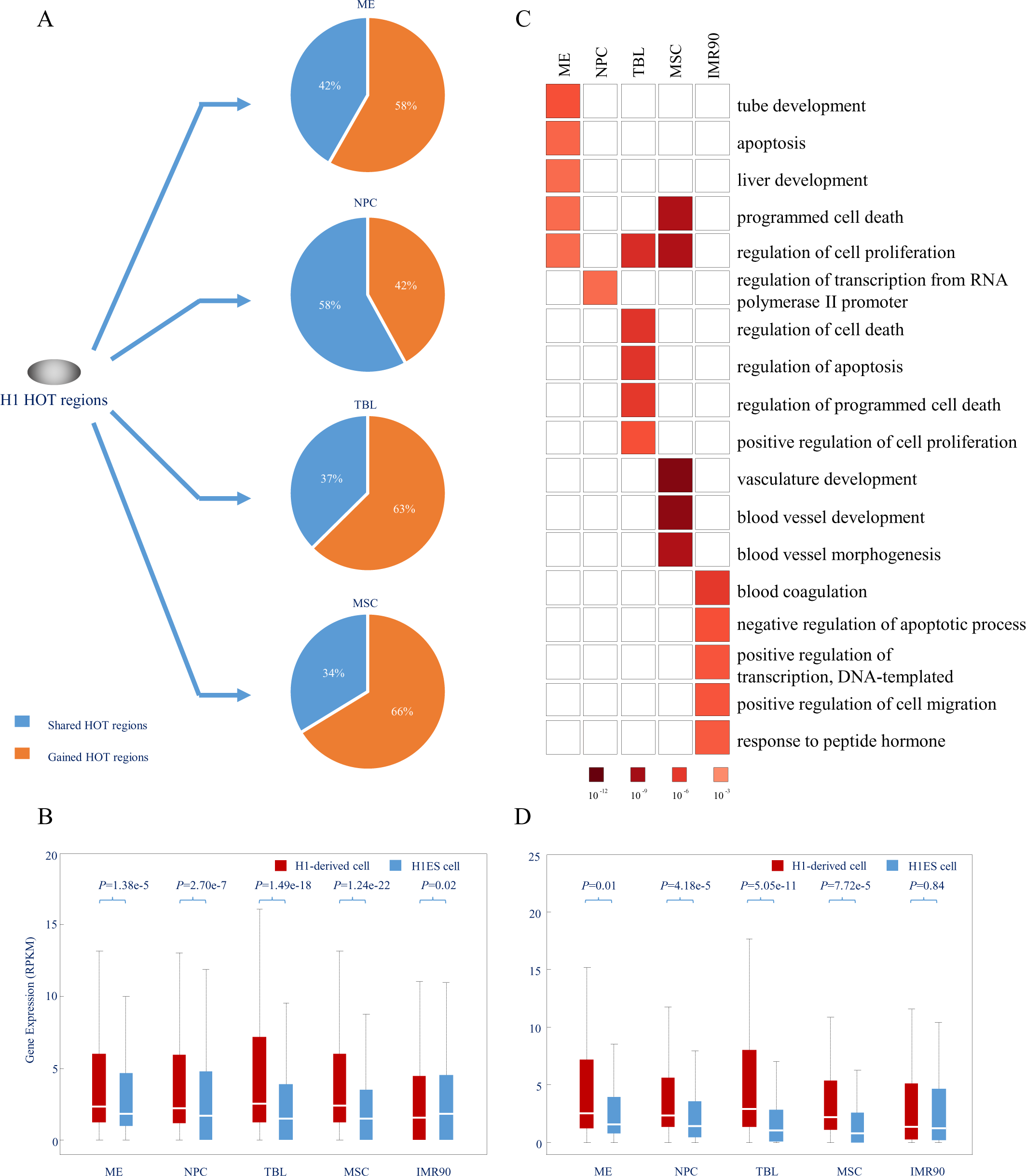
Dynamics of HOT regions during H1 differentiation. (A) Dynamic changes in HOT regions during the differentiation of H1 cell to its derived cells. The proportions of “gained” HOT regions (orange) and “shared” HOT regions (blue) in H1-derived cells are shown. (B) Boxplot showing the expression of genes associated with “gained” HOT regions in H1-derived cells and the IMR90 cell type (red) and in H1ES cells (blue), *p*-values were calculated by the Kolmogorov-Smirnov test. (C) GO analysis of genes associated with “gained” HOT regions in H1-derived cells and the IMR90 cell type using DAVID. For each cell type, top five biological processes were shown. (D) Boxplot showing the expression of enriched TF genes in HOT regions of H1-derived cells and the IMR90 cell type (red) and in H1ES cells (blue); *p*-values were calculated by the Kolmogorov-Smirnov test. See also Table S8-S9.

Then, we scanned TF motifs in HOT regions of H1 and its derived cell types using iFORM [23] and identified 106, 153, 178, 75 and 35 enriched TF genes in ME, NPC, TBL, MSC and IMR90 cells, respectively (Materials and Methods, Table S9). These enriched TF genes were highly expressed in H1-derived cell types relative to H1 cells (*p*-value < 0.01, Kolmogorov-Smirnov test), respectively (Fig. 5D). Further GO analysis demonstrated that these enriched TF genes play key roles in the regulation of transcription, gene expression, and metabolic process during the differentiation of H1 into its derived cell types (Table S9). Taken together, our results improve our understanding of HOT region dynamics during development and differentiation.

### Epigenetic regulation of HOT regions upon H1 differentiation

To further investigate the epigenetic regulatory mechanism of HOT regions upon development and differentiation, we examined the potential role of histone modifications at HOT regions upon H1 differentiation.

Previously, bivalent genes marked by both H3K4me3 and H3K27me3 were shown to be highly associated with developmental genes [36]. Intriguingly, an analysis of the data from previous studies [37, 38] showed that the genes associated with HOT regions during H1 differentiation appeared to be highly enriched in bivalent genes (*p*-value < 1.77×10^−9^, hypergeometric test, Table S10). We then asked whether HOT regions undergo dynamic epigenetic regulation upon development and differentiation. We examined the dynamic epigenetic modifications at these HOT regions upon H1 differentiation and found that over 50% of HOT regions in H1 cells were bound simultaneously by H3K4me3 and H3K27me3. We named these regions bivalent HOT regions. Interestingly, the number of bivalent HOT regions decreased upon H1 differentiation. In differentiated cells, a large portion of HOT regions were bound only by H3K4me3 or H3K27me3 relative to H1 cells (Fig. 6A). This is in good agreement with the opinion that bivalent genes become monovalent upon cell differentiation [36]. Furthermore, over three-quarters of bivalent HOT regions in H1 cells were “lost”, whereas less than one-quarter of bivalent HOT regions in H1 cells were “shared” during the differentiation of H1 into its derived cell types (Fig. 6B). Moreover, a considerable proportion (ranging from 10% to 38%) of “lost” bivalent regions were differentiated into “activated” HOT regions bound by H3K4me3 only, whereas only a very small proportion of these regions differentiated into HOT regions bound only by H3K27me3 alone (Fig. 6B).

**Figure 6.** Epigenetic Regulation of HOT regions upon H1 differentiation. (A) The chromatin state (presence of H3K4me3 and/or H3K27me3) of HOT regions in various cell types. (B) Dynamic changes of bivalent HOT regions between H1 to H1-derived cell types. (C) Box plots showing the levels of H3k4me3 (top), H3k27me3 (middle), and mRNA (bottom) at activated and repressive HOT regions in H1 and H1-derived cells. See also Table S10-S11.

We further explored the dynamic epigenetic signals upon H1 differentiation. We found that in H1-derived cells, the “activated” HOT regions were marked by a higher level of H3k4me3 and a lower level of H3K27me3 compared to the H1 bivalent HOT regions. Moreover, the “repressed” HOT regions in H1-derived cells were marked by a lower level of H3k4me3 and a higher level of H3K27me3 compared to the H1 bivalent HOT regions. The levels of H3K4me3 and H3K27me3 in the activated HOT regions of H1-derived cells increased and decreased during differentiation, respectively. A gene expression analysis of the “activated” and “repressed” HOT regions further confirmed these findings (Fig. 6C). Furthermore, we performed GO analysis of the bivalent HOT regions in H1 and the activated HOT regions in H1-derived cells and found that the genes associated with these regions were strongly enriched for the functional categories “regulation of transcription”, “metabolic process” and “differentiation” (Table S11). Taken together, our findings reveal a preliminary view of the dynamic epigenetic regulation of HOT regions, which were strongly associated with developmental genes and had key roles upon cell differentiation.

## Discussion and Conclusions

Previous studies have revealed regions in worms [1, 2], flies [3–7], and humans [8–10] with heavily clustered TF binding that have been termed HOT regions. These reports [1–10] identified HOT regions by the binding peaks of many TFs using ChIP-seq data, whereas we defined HOT regions by a large number of TF motif binding sites on DHSs in DNase-seq data. This change was made because previous investigations of experimental HOT regions were restricted by the currently limited amount of TF ChIP-seq data and by the consistency of HOT regions defined by identical combinations of TFs across diverse cells/tissues. Although the identifications of HOT regions were based on different data, both definitions demonstrate that HOT regions are a novel class of genomic regions that are bound by a surprisingly large number of TFs and contain numerous TF motifs. Importantly, our identification of HOT regions using TF motif discovery on DHSs can greatly extend the repertoire of both TFs and cell types in the genome, thus greatly enhancing our understanding of HOT regions.

We have extended our understanding of HOT regions by demonstrating that ESC HOT regions are highly transcribed and by identifying the population of TFs, cofactors, chromatin regulators, and core transcription machinery that occupy these domains in ESCs. ESCs were chosen for identifying components of HOT regions because the TFs, cofactors, chromatin regulators, and noncoding RNAs that control the ESC state and that contribute to the gene expression programmes of pluripotency and self-renewal are likely better understood than those for any other cell type [39–41]. HOT regions are occupied by a large portion of enhancer-associated RNA polymerase II and its associated cofactors and chromatin regulators, which may explain how these molecules contribute to the high transcriptional levels of genes associated with HOT regions. Furthermore, the levels of RNA detected in HOT regions vastly exceed the levels of RNA in LOT regions, and recent evidence suggests that these enhancer RNAs (eRNAs) may contribute to gene activation [42–49]. Several additional important insights were gained by studying how more than 40 TFs, cofactors, chromatin regulators, and components of the core transcription machinery occupy HOT regions and LOT regions in ESCs. All of the enhancer-binding TFs are enriched in HOT regions, with some so highly enriched that they distinguish HOT regions from LOT regions.

By uncovering characteristic sequence signatures of HOT regions, our computational analysis revealed that more than one quarter of enriched TF motifs exhibited significantly enriched binding within HOT regions; the majority of these TF motifs play essential roles in development and cell differentiation. Strikingly, 12 of 34 TFs (*p*-value = 0.0012, binomial test) that showed specifically enriched binding within LOT regions were housekeeping TFs. In combination with previous observations that HOT regions are depleted in the bound TFs’ motifs [1, 3–5] compared with regions bound by single TFs, our findings suggest that HOT regions have distinct sequence features that distinguish them from LOT regions and the genome background. Moreover, these findings suggest that information regarding HOT regions is encoded in the DNA sequence.

We have generated a catalogue of HOT regions and their associated genes in a broad spectrum of human cell and tissue types. HOT regions tend to be cell type-specific, and the genes associated with these elements are linked to biological processes that largely define the development and differentiation of the respective cell and tissue types. Genes that encode candidate key developmental TFs and noncoding RNAs, such as microRNAs, are among those associated with HOT regions. Thus, the HOT region catalogue should be a valuable resource for further studies of the transcriptional control of cell development and differentiation [50–53]. Based on the catalogue of HOT regions, our further study has explored the association of GWAS SNPs and HOT regions, and our findings have illustrated the key roles of HOT regions in human disease and cancer [54].

An association analysis between HOT regions and embryonic enhancers presented direct evidence of the systematic developmental persistence of tissue-selective early developmental enhancers at HOT regions and of the persistent imprint of enhancer roles on the formation of cross-cell-type patterning of HOT regions in definitive cells. Additionally, we found that super-enhancers were highly enriched in HOT regions across diverse cell types, and cell type-specific super-enhancers tend to strongly colocalise with the HOT regions that function in the corresponding cell types. Furthermore, all enhancer markers, including DNaseI, H3K27ac, H3K4me1, enhancer-binding TFs and chromatin regulators, are enriched at HOT regions but have lower levels of enrichment that distinguish them from super-enhancers. Strikingly, we observed the paradoxical coexistence of permissive and repressive histone marks, H3K4me3 and H3K27me3, in HOT regions. Although GO analysis revealed that super-enhancers drive the expression of genes that define the identity of the respective cell and tissue types, HOT regions are associated with biological processes that largely define the development and differentiation of the respective cell and tissue types. Together, our results suggest that HOT regions might therefore represent a novel class of enhancers because they contain many discriminatory features that are different from enhancers or super-enhancers. The activities of HOT regions and super-enhancers are both defined by the colocalisation of TFs in these regions but on different genomic scales of colocalisation. A recent study [55] described the relationship between hotspots and super-enhancers in the early phase of adipogenesis, demonstrating that hotspots are highly enriched in large super-enhancers and revealing that hotspots and super-enhancers function as two levels of regulatory hubs that serve to integrate external stimuli through cooperativity between TFs on chromatin. These findings are highly consistent with ours.

Finally, we examined the dynamics changes in HOT regions and correlated their associated genes and enriched TF genes during the differentiation of H1 into its derived cell types. We observed that these associated genes and enriched TF genes were highly expressed in H1-derived cell types relative to H1 cells and play key roles during development or differentiation. We further examined the dynamic epigenetic regulation at HOT regions during H1 differentiation. We found that a large proportion of HOT regions in H1 cells demonstrated a bivalent state, and the portion of the bivalent HOT regions decreased during the differentiation of the hESC line H1 into ME cells, TBL cells, NPCs, and MSCs. Many of these bivalent HOT regions were differentiated into activated regions. Additionally, we demonstrated that the activated regions showed an inverse relationship with the levels of H3k4me3 and H3k27me3. Our results present a preliminary view of the dynamic regulation of HOT regions upon cell differentiation and will improve our understanding of HOT region dynamics during development and differentiation.

Taken together, our findings provide a resource for the functional exploration of HOT regions and extend our understanding of the key roles of HOT regions during development and differentiation.

## Materials and Methods

### Data sets

The DNaseI Hypersensitivity by Digital DNaseI data were obtained from the Duke and UW ENCODE groups. Histone modifications according to ChIP-seq data were downloaded from the Broad histone ENCODE group. TFs according to ChIP-seq data were obtained from the HAIB and SYDH TFBS ENCODE groups. DNase-seq and ChIP-seq data in both peak file and bam file formats were used in this study. Gene annotations were obtained from the GENCODE data (V15) [56]. All these data were collected from the ENCODE Project [24], and the use of these data strictly adhered to the ENCODE Consortium Data Release Policy. The data used for epigenomic analysis of HOT regions in H1 cells and four H1-derived lineages were obtained from a recent study [29].

### Identifying TFBS-clustered Regions and HOT regions

Position weight matrices (PWMs) of 542 TFs, which corresponded to 796 motif models, were collected from the TRANSFAC [57], JASPAR [58], and UniPROBE [59] databases. The genomic sequence under DHSs from the hg19 genome was used as the input for iFORM [23] using a custom library of all 796 motifs scanned for motif instances at a *p*-value threshold of 10^−18^ (corresponding to the FIMO threshold of 10^−5^). For each TF, instances of multiple TF motifs were combined to generate the corresponding TFBSs.

An established method [5] was used to perform Gaussian kernel density estimations across the genome (bandwidth 3 kb, centred on each TFBS). Each peak of the density profile was denoted as a TFBS-clustered region. To determine the complexity of each TFBS-clustered region, the Gaussian kernalised distance from a peak to each TFBS that contributed at least 0.1 to the strength was determined. The window around each TFBS-clustered region was derived by finding the maximum distance (in bp) from the TFBS-clustered region to a contributing TF and adding 1.5 kb (one half of the bandwidth). Each window was centred on a TFBS-clustered region.

To identify HOT regions, we first ranked all the TFBS-clustered regions in a cell type by increasing and plotting the TFBS complexity (Fig. 1A). This plot revealed a clear point in the distribution of the TFBS-clustered regions at which the complexity signal began to increase rapidly. To geometrically define this point, we first scaled the data such that the *x* and *y* axes were from 0-1. We then found the x axis point for which a line with a slope of 1 was tangent to the curve. We defined the TFBS-clustered regions above this point to be HOT regions and the TFBS-clustered regions below that point to be LOT regions. The pipeline for identifying HOT or LOT regions was applied uniformly to datasets from 349 samples, including 154 cell types studied under the ENCODE Project [24]. The classification of the TFBS-clustered regions as a HOT or LOT region in each cell type for diverse human cells and tissues can be found in Table S2.

### Validation of HOT regions with ChIP-seq

We downloaded publicly available HOT regions defined based on the ChIP-seq data from the ENCODE Consortium obtained in five cell types, including K562, Hep-G2, HeLa-S3, H1-hESC, and GM12878 cells [24, 25]. First, we used GSC (genome structure correction) analysis to assess the performance of predicting HOT regions. The GSC statistic [60, 61] was used to calculate the confidence intervals (CIs) for the experimental HOT regions that were expected to contain our HOT regions by chance. This statistic provides a conservative correction to standard tests. Additionally, we applied the ROC (receiver operating characteristic) curves and the corresponding area under the curve (AUC) to validate our HOT regions with the experimental HOT and LOT regions in the five cell types as the “gold-standard” data sets.

To further verify whether TFs indeed bound within the identified HOT regions, we collected uniform ChIP-seq peaks corresponding to multiple TFs from the ENCODE Project in the five cell types. We counted the occurrence rates of ChIP-seq peaks of diverse TFs that were located within our HOT regions and experimental HOT regions. Additionally, we explored the correlations between the TFBS complexity of our HOT regions and the number of TF peaks that were located within our HOT regions.

### Identifying motifless binding peaks

For each TF, we collected the ChIP-seq binding peaks in each cell line and scanned the TF motifs of each binding peak using iFORM [23]. We identified the binding peaks that do not have DNA sequence motifs of the corresponding TFs using the method presented in a previous study [25]. Then, we collected the peaks of all TFs to construct the set of motifless binding peaks for each cell line.

We then associated the motifless binding peaks with our HOT regions and experimental HOT regions, respectively. Because our HOT regions and experimental HOT regions were identified in the whole human genome but the motifless binding peaks were defined from ChIP-seq binding peaks, we first identified the subsets of our HOT regions and experimental HOT regions located within these ChIP-seq binding peaks for each cell line. We then determined the association of the motifless binding regions with the subsets of HOT regions and used GSC analysis to measure the statistical significance of these associations.

### Master list and annotation for HOT regions

The HOT regions from 154 cell types were consolidated into a master list of 59,986 unique, non-overlapping HOT region positions by first merging these regions across cell types. Then, for each resulting interval of merged regions, the HOT region with the highest TFBS complexity was selected for the master list. Any HOT regions overlapping the regions selected for the master list were then discarded. The remaining HOT regions were merged, and the process was repeated until each original TFBS-clustered region was either incorporated into the master list or discarded.

Genomic annotations from GENCODE annotations (V15) [56], i.e., Basic, Comprehensive, PseudoGenes, 2-way PseudoGenes, and PolyA Transcripts, were used. The promoter (proximal) class of each GENCODE-annotated TSS was defined as a region from the master list within 1 kb of the TSS. The exon class was defined as any HOT region not in the promoter class that overlapped a GENCODE-annotated “CDS” segment by at least 75 bp. The UTR class was defined as a HOT region not in the promoter or exon class that overlapped a GENCODE-annotated “UTR” segment by at least 1 bp. The intron class was defined as a GENCODE segment annotated as “gene” with all “CDS” segments. The intron class also covered any HOT regions not defined by other categories that overlapped introns by at least 1 bp.

The distal category was defined as the HOT regions located at least 2 kb away from any GENCODE-annotated TSS. Repeat categories for the LINE, SINE, LTR, and DNA repeat classes were obtained from the UCSC RepeatMasker track annotations. The miRNA category counts for each miRNA annotated by miRBase (version 20) [26, 62] were defined by the closest master list HOT regions within 1 kb upstream and downstream of the miRNA TSS.

The cell-type number was defined for each HOT region by annotating the master list with the number of cell types with overlapping HOT regions. The plots in Figure 1E were generated using the R function “geom_violin” from the “ggplot2” package, which summarises the distribution of cell type numbers for distinct categories of HOT regions. The distribution of cell types containing a HOT region was calculated separately for HOT regions observed in 154 cell types.

### Association analysis of functional regulatory elements

The microRNA coordinates were downloaded from miRBase (version 20) [26] and used to map microRNAs to their genomic locations. We used the method described in a recent study [15] to assign TSSs for 2633 microRNA loci. RepeatMasker data were downloaded from the hg19 rmsk table associated with the UCSC Genome Browser. There are 1395 distinctly named repeats in 56 families in 21 repeat classes. The data were analysed by repeat family because this procedure gives a granularity suitable for display. A number of the classes are structural classes rather than classes derived from transposable elements. Bedops utilities [63] were used to count the number of repeat elements that overlapped at least 1 bp with HOT regions. The HOT regions from 154 cell types/tissues were tested for overlap with repeat families. Supplementary Table S5 shows overlap statistics for families of elements with at least 5000 overlapping HOT regions. Additionally, an extensive compilation of 373 experimentally validated distal, non-promoter *cis*-regulatory elements, including insulators, locus control regions, and so on, were taken from a recent study [15] (Table S6). Finally, we collected low methylation regions (LMRs), unmethylated regions (UMRs), and DNA methylation valleys (DMVs) from a recent study [29].

### Comparison of HOT regions with known enhancers

We downloaded the data for tests of human enhancers in a mouse developmental model [30, 64] from the VISTA enhancer browser http://enhancer.lbl.gov/ and received permission to use the embryonic mouse images.

To calculate the enrichment of super-enhancers in a HOT region, we performed a GSC analysis between HOT regions and super-enhancers, as shown in Fig. 4A. To perform the colocalisation analysis on HOT regions and super-enhancers in *N*-by-*N* (*N* = 14) cell line combinations as similarly described in a previous study [34, 35], we collected a catalogue of super-enhancers in 14 human cell types from recent studies.

The counts were divided by the corresponding row sum and column sum and multiplied by the matrix sum to obtain enrichment values using the same approach as the *χ*^2^ test. We plotted the enrichment factor for each histone modification in an *N*-by-*N* heat map.

### Dynamics of HOT regions upon H1 differentiation

Using the DNase-seq data obtained from a recent study [29], we identified 5,410, 5,602, 4,250, 3,408, and 6,719 HOT regions in H1-hESCs (H1), mesendoderm (ME), neural progenitor cells (NPCs), trophoblast-like cells (TBL), and mesenchymal stem cells (MSCs), respectively. As a control for terminally differentiated cells, we also identified 4,002 HOT regions in IMR90, a primary human foetal lung fibroblast cell line. Each HOT region was assigned to the closest genes annotated in GENCODE (V15) [56] by determining the distance from the centre of the HOT region to the TSS of each GENCODE gene.

HOT regions that were “shared” upon H1 differentiation were defined as HOT regions belonging to both H1 cells and H1-derived cell types (using *bedops –e −25%*). HOT regions “lost” upon H1 differentiation were defined as HOT regions belonging to H1 cells but not found within H1-derived cells (using *bedops –n −25%*). HOT regions that were “gained” upon H1 differentiation were defined as regions belonging to H1-derived cells but not found within H1 cells (using *bedops –n −25%*). The genes associated with “gained” HOT regions were further restricted to those that expressed (RPKM > 1) in either H1 or its derived cells.

We scanned TF motifs in HOT regions of H1 and H1-derived cells using iFORM [23]. We calculated the TF density of each cell and took the ratio of densities between H1 and its derived cells. We defined as enriched TF gene those TF genes with ratio > 1 and expressed > 1 (RPKM > 1) in either H1 cells or its derived cells.

MACS (Zhang et al., 2008) was used to identify H3k4me3 peaks using default parameters. For H3k27me3, which typically exhibits broad enrichment, we used RSEG [65] to identify enriched regions with “*−i 20 −b 100 −v −mode 2*” Upon H1 differentiation, “activated” HOT regions were defined as HOT regions bearing H3k4me3 peaks only, whereas “repressed” HOT regions were defined as HOT regions bearing either H3k27me3 peaks only or no marker.

We used DAVID [66] to perform GO analysis of genes associated with “gained” HOT regions, TF-enriched genes, and genes associated with “activated” HOT regions.

## Accession numbers

The identified HOT regions across human cell and tissue types have been deposited in the Gene Expression Omnibus under the accession ID GSE54296.

## Acknowledgements

We wish to thank the ENCODE Project Consortium for making their data publicly available. This work was supported by grants from the Major Research plan of the National Natural Science Foundation of China (No. U1435222), the Program of International S&T Cooperation (No. 2014DFB30020) and the National High Technology Research and Development Program of China (No. 2015AA020108).

## Author Contributions

W.S. conceived the project. W.S., X.B. and S.W. designed all experiments. H.L., H.C., and F.L. performed the experiments. All authors analysed the data and contributed to the manuscript preparation. W.S. wrote the manuscript.

## Additional information

Competing financial interests: The authors declare no competing financial interests.

## References

1 Gerstein MB, Lu ZJ, Van Nostrand EL, Cheng C, Arshinoff Bl, Liu T, Yip KY, Robilotto R, Rechtsteiner A, Ikegami K et al: Integrative analysis of the Caenorhabditis elegans genome by the modENCODE project. Science (New York, NY) 2010, 330(6012): 1775–1787.

2 Araya CL, Kawli T, Kundaje A, Jiang L, Wu B, Vafeados D, Terrell R, Weissdepp P, Gevirtzman L, Mace D et al: Regulatory analysis of the C. elegans genome with spatiotemporal resolution. Nature 2014, 512(7515): 400–405.

3 Moorman C, Sun LV, Wang J, de Wit E, Talhout W, Ward LD, Greil F, Lu XJ, White KP, Bussemaker HJ et al: Hotspots of transcription factor colocalization in the genome of Drosophila melanogaster. Proceedings of the National Academy of Sciences of the United States of America 2006, 103(32): 12027–12032.

4 MacArthur S, Li XY, Li J, Brown JB, Chu HC, Zeng L, Grondona BP, Hechmer A, Simirenko L, Keranen SV et al: Developmental roles of 21 Drosophila transcription factors are determined by quantitative differences in binding to an overlapping set of thousands of genomic regions. Genome biology 2009, 10(7):R80.

5 Roy S, Ernst J, Kharchenko PV, Kheradpour P, Negre N, Eaton ML, Landolin JM, Bristow CA, Ma L, Lin MF et al: Identification of functional elements and regulatory circuits by Drosophila modENCODE. Science (New York, NY) 2010, 330(6012): 1787–1797.

6 Negre N, Brown CD, Ma L, Bristow CA, Miller SW, Wagner U, Kheradpour P, Eaton ML, Loriaux P, Sealfon R et al: A cis-regulatory map of the Drosophila genome. Nature 2011, 471(7339): 527–531.

7 Kvon EZ, Stampfel G, Yanez-Cuna JO, Dickson BJ, Stark A: HOT regions function as patterned developmental enhancers and have a distinct cis-regulatory signature. Genes & development 2012, 26(9): 908–913.

8 Yan J, Enge M, Whitington T, Dave K, Liu J, Sur I, Schmierer B, Jolma A, Kivioja T, Taipale M et al: Transcription factor binding in human cells occurs in dense clusters formed around cohesin anchor sites. Cell 2013, 154(4): 801–813.

9 Chen RA, Stempor P, Down TA, Zeiser E, Feuer SK, Ahringer J: Extreme HOT regions are CpG-dense promoters in C. elegans and humans. Genome research 2014, 24(7): 1138–1146.

10 Foley JW, Sidow A: Transcription-factor occupancy at HOT regions quantitatively predicts RNA polymerase recruitment in five human cell lines. BMC genomics 2013, 14:720.

11 Furlong EE: Molecular biology: A fly in the face of genomics. Nature 2011, 471(7339): 458–459.

12 Blaxter M: Genetics. Revealing the dark matter of the genome. Science (New York, NY) 2010, 330(6012): 1758–1759.

13 Grant CE, Bailey TL, Noble WS: FIMO: scanning for occurrences of a given motif. Bioinformatics (Oxford, England) 2011, 27(7): 1017–1018.

14 Heinz S, Benner C, Spann N, Bertolino E, Lin YC, Laslo P, Cheng JX, Murre C, Singh H, Glass CK: Simple combinations of lineage-determining transcription factors prime cis-regulatory elements required for macrophage and B cell identities. Molecular cell 2010, 38(4): 576–589.

15 Thurman RE, Rynes E, Humbert R, Vierstra J, Maurano MT, Haugen E, Sheffield NC, Stergachis AB, Wang H, Vernot B et al: The accessible chromatin landscape of the human genome. Nature 2012, 489(7414): 75–82.

16 Gaszner M, Felsenfeld G: Insulators: exploiting transcriptional and epigenetic mechanisms. Nature reviews Genetics 2006, 7(9): 703–713.

17 Gross DS, Garrard WT: Nuclease hypersensitive sites in chromatin. Annual review of biochemistry 1988, 57: 159–197.

18 Li Q, Harju S, Peterson KR: Locus control regions: coming of age at a decade plus. Trends in genetics 1999, 15(10): 403–408.

19 Boyle AP, Davis S, Shulha HP, Meltzer P, Margulies EH, Weng Z, Furey TS, Crawford GE: High-resolution mapping and characterization of open chromatin across the genome. Cell 2008, 132(2)311–322.

20 Hesselberth JR, Chen X, Zhang Z, Sabo PJ, Sandstrom R, Reynolds AP, Thurman RE, Neph S, Kuehn MS, Noble WS et al: Global mapping of protein-DNA interactions in vivo by digital genomic footprinting. Nature methods 2009, 6(4): 283–289.

21 John S, Sabo PJ, Thurman RE, Sung MH, Biddie SC, Johnson TA, Hager GL, Stamatoyannopoulos JA: Chromatin accessibility pre-determines glucocorticoid receptor binding patterns. Nature genetics 2011, 43(3): 264–268.

22 Chen H, Li H, Liu F, Zheng X, Wang S, Bo X, Shu W: An integrative analysis of TFBS-clustered regions reveals new transcriptional regulation models on the accessible chromatin landscape. Scientific reports 2015, 5: 8465.

23 Ren C, Chen H, Liu F, Li H, Bo X, Shu W: iFORM: incorporating Find Occurrence of Regulatory Motifs. bioRxiv 2016.

24 Consortium TEP: An integrated encyclopedia of DNA elements in the human genome. Nature 2012, 489(7414):57–74.

25 Yip KY, Cheng C, Bhardwaj N, Brown JB, Leng J, Kundaje A, Rozowsky J, Birney E, Bickel P, Snyder M et al: Classification of human genomic regions based on experimentally determined binding sites of more than 100 transcription-related factors. Genome biology 2012, 13(9):R48.

26 Kozomara A, Griffiths-Jones S: miRBase: annotating high confidence microRNAs using deep sequencing data. Nucleic acids research 2014, 42(Database issue):D68–73.

27 Djebali S, Davis CA, Merkel A, Dobin A, Lassmann T, Mortazavi A, Tanzer A, Lagarde J, Lin W, Schlesinger F et al: Landscape of transcription in human cells. Nature 2012, 489(7414): 101–108.

28 Stadler MB, Murr R, Burger L, Ivanek R, Lienert F, Scholer A, van Nimwegen E, Wirbelauer C, Oakeley EJ, Gaidatzis D et al: DNA-binding factors shape the mouse methylome at distal regulatory regions. Nature 2011, 480(7378): 490–495.

29 Xie W, Schultz MD, Lister R, Hou Z, Rajagopal N, Ray P, Whitaker JW, Tian S, Hawkins RD, Leung D et al: Epigenomic analysis of multilineage differentiation of human embryonic stem cells. Cell 2013, 153(5): 1134–1148.

30 Pennacchio LA, Ahituv N, Moses AM, Prabhakar S, Nobrega MA, Shoukry M, Minovitsky S, Dubchak I, Holt A, Lewis KD et al: In vivo enhancer analysis of human conserved non-coding sequences. Nature 2006, 444(7118): 499–502.

31 Hnisz D, Abraham BJ, Lee TI, Lau A, Saint-Andre V, Sigova AA, Hoke HA, Young RA: Super-enhancers in the control of cell identity and disease. Cell 2013, 155(4): 934–947.

32 Loven J, Hoke HA, Lin CY, Lau A, Orlando DA, Vakoc CR, Bradner JE, Lee TI, Young RA: Selective inhibition of tumor oncogenes by disruption of super-enhancers. Cell 2013, 153(2)320–334.

33 Whyte WA, Orlando DA, Hnisz D, Abraham BJ, Lin CY, Kagey MH, Rahl PB, Lee TI, Young RA: Master transcription factors and mediator establish super-enhancers at key cell identity genes. Cell 2013, 153(2)307–319.

34 Xi H, Shulha HP, Lin JM, Vales TR, Fu Y, Bodine DM, McKay RD, Chenoweth JG, Tesar PJ, Furey TS et al: Identification and characterization of cell type-specific and ubiquitous chromatin regulatory structures in the human genome. PLoS genetics 2007, 3(8):el36.

35 Chen H, Tian Y, Shu W, Bo X, Wang S: Comprehensive identification and annotation of cell type-specific and ubiquitous CTCF-binding sites in the human genome. PloS one 2012, 7(7):e41374.

36 Bernstein BE, Mikkelsen TS, Xie X, Kamal M, Huebert DJ, Cuff J, Fry B, Meissner A, Wernig M, Plath K et al: A bivalent chromatin structure marks key developmental genes in embryonic stem cells. Cell 2006, 125(2)315–326.

37 Zhao XD, Han X, Chew JL, Liu J, Chiu KP, Choo A, Orlov YL, Sung WK, Shahab A, Kuznetsov VA et al: Whole-genome mapping of histone H3 Lys4 and 27 trimethylations reveals distinct genomic compartments in human embryonic stem cells. Cell stem cell 2007, 1(3):286–298.

38 Pan G, Tian S, Nie J, Yang C, Ruotti V, Wei H, Jonsdottir GA, Stewart R, Thomson JA: Whole-genome analysis of histone H3 lysine 4 and lysine 27 methylation in human embryonic stem cells. Cell stem cell 2007, 1(3): 299–312.

39 Ng HH, Surani MA: The transcriptional and signalling networks of pluripotency. Nature cell biology 2011, 13(5): 490–496.

40 Orkin SH, Hochedlinger K: Chromatin connections to pluripotency and cellular reprogramming. Ce// 2011, 145(6)335–850.

41 Young RA: Control of the embryonic stem cell state. Cell 2011, 144(6): 940–954.

42 Kim TK, Hemberg M, Gray JM, Costa AM, Bear DM, Wu J, Harmin DA, Laptewicz M, Barbara-Haley K, Kuersten S et al: Widespread transcription at neuronal activity-regulated enhancers. Nature 2010, 465(7295): 182–187.

43 Lai F, Orom UA, Cesaroni M, Beringer M, Taatjes DJ, Blobel GA, Shiekhattar R: Activating RNAs associate with Mediator to enhance chromatin architecture and transcription. Nature 2013, 494(7438):497–501.

44 Orom UA, Derrien T, Beringer M, Gumireddy K, Gardini A, Bussotti G, Lai F, Zytnicki M, Notredame C, Huang Q et al: Long noncoding RNAs with enhancer-like function in human cells. Cell 2010, 143(1):46–58.

45 Ling J, Ainol L, Zhang L, Yu X, Pi W, Tuan D: HS2 enhancer function is blocked by a transcriptional terminator inserted between the enhancer and the promoter. The Journal of biological chemistry 2004, 279(49): 51704–51713.

46 Kaikkonen MU, Spann NJ, Heinz S, Romanoski CE, Allison KA, Stender JD, Chun HB, Tough DF, Prinjha RK, Benner C et al: Remodeling of the enhancer landscape during macrophage activation is coupled to enhancer transcription. Molecular cell 2013, 51(3)310–325.

47 Mousavi K, Zare H, Dell’orso S, Grontved L, Gutierrez-Cruz G, Derfoul A, Hager GL, Sartorelli V: eRNAs promote transcription by establishing chromatin accessibility at defined genomic loci. Molecular cell 2013, 51(5): 606–617.

48 Lam MT, Cho H, Lesch HP, Gosselin D, Heinz S, Tanaka-Oishi Y, Benner C, Kaikkonen MU, Kim AS, Kosaka M et al: Rev-Erbs repress macrophage gene expression by inhibiting enhancer-directed transcription. Nature 2013, 498(7455): 511–515.

49 Li W, Notani D, Ma Q, Tanasa B, Nunez E, Chen AY, Merkurjev D, Zhang J, Ohgi K, Song X et al: Functional roles of enhancer RNAs for oestrogen-dependent transcriptional activation. Nature 2013, 498(7455): 516–520.

50 Zhou Q, Brown J, Kanarek A, Rajagopal J, Melton DA: In vivo reprogramming of adult pancreatic exocrine cells to beta-cells. Nature 2008, 455(7213): 627–632.

51 Lee Tl, Young RA: Transcriptional regulation and its misregulation in disease. Cell 2013, 152(6): 1237–1251.

52 Graf T, Enver T: Forcing cells to change lineages. Nature 2009, 462(7273): 587–594.

53 Cherry AB, Daley GQ: Reprogramming cellular identity for regenerative medicine. Cell 2012, 148(6): 1110–1122.

54 Li H, Chen H, Liu F, Ren C, Wang S, Bo X, Shu W: Functional annotation of HOT regions in the human genome: implications for human disease and cancer. Scientific reports 2015, 5:11633.

55 Siersbaek R, Rabiee A, Nielsen R, Sidoli S, Traynor S, Loft A, La Cour Poulsen L, Rogowska-Wrzesinska A, Jensen ON, Mandrup S: Transcription factor cooperativity in early adipogenic hotspots and super-enhancers. Cell reports 2014, 7(5): 1443–1455.

56 Harrow J, Frankish A, Gonzalez JM, Tapanari E, Diekhans M, Kokocinski F, Aken BL, Barrell D, Zadissa A, Searle S et al: GENCODE: the reference human genome annotation for The ENCODE Project. Genome research 2012, 22(9): 1760–1774.

57 Matys V, Kel-Margoulis OV, Fricke E, Liebich I, Land S, Barre-Dirrie A, Reuter I, Chekmenev D, Krull M, Hornischer K et al: TRANSFAC and its module TRANSCompel: transcriptional gene regulation in eukaryotes. Nucleic acids research 2006, 34(Database issue):D108–110.

58 Portales-Casamar E, Thongjuea S, Kwon AT, Arenillas D, Zhao X, Valen E, Yusuf D, Lenhard B, Wasserman WW, Sandelin A: JASPAR 2010: the greatly expanded open-access database of transcription factor binding profiles. Nucleic acids research 2010, 38(Database issue):D105–110.

59 Robasky K, Bulyk ML: UniPROBE, update 2011: expanded content and search tools in the online database of protein-binding microarray data on protein-DNA interactions. Nucleic acids research 2011, 39(Database issue):D124–128.

60 Bickel PJ, Boley N, Brown JB, Huang H, Zhang NR: Subsampling methods for genomic inference. 2010: 1660–1697.

61 Birney E, Stamatoyannopoulos JA, Dutta A, Guigo R, Gingeras TR, Margulies EH, Weng Z, Snyder M, Dermitzakis ET, Thurman RE et al: Identification and analysis of functional elements in 1% of the human genome by the ENCODE pilot project. Nature 2007, 447(7146): 799–816.

62 Griffiths-Jones S, Saini HK, van Dongen S, Enright AJ: miRBase: tools for microRNA genomics. Nucleic acids research 2008, 36(Database issue):D154–158.

63 Neph S, Kuehn MS, Reynolds AP, Haugen E, Thurman RE, Johnson AK, Rynes E, Maurano MT, Vierstra J, Thomas S et al: BEDOPS: high-performance genomic feature operations. Bioinformatics (Oxford, England) 2012, 28(14): 1919–1920.

64 May D, Blow MJ, Kaplan T, McCulley DJ, Jensen BC, Akiyama JA, Holt A, Plajzer-Frick I, Shoukry M, Wright C et al. Large-scale discovery of enhancers from human heart tissue. Nature genetics 2012, 44(1):89–93.

65 Song Q, Smith AD: Identifying dispersed epigenomic domains from ChIP-Seq data. Bioinformatics (Oxford, England) 2011, 27(6): 870–871.

66 Huang da W, Sherman BT, Lempicki RA: Systematic and integrative analysis of large gene lists using DAVID bioinformatics resources. Nature protocols 2009, 4(1):44–57.

